# Deciphering the Differential Impact of Thrombopoietin/MPL Signaling on Hematopoietic Stem Cell Function in Bone Marrow and Spleen

**DOI:** 10.1101/2023.04.27.538580

**Authors:** Sandy Lee, Huichun Zhan

## Abstract

Thrombopoietin (TPO) and its receptor MPL play crucial roles in hematopoietic stem cell (HSC) function and platelet production. However, the precise effects of TPO/MPL signaling on HSC regulation in different hematopoietic niches remain unclear. Here, we investigated the effects of TPO/MPL ablation on marrow and splenic hematopoiesis in TPO^-/-^ and MPL^-/-^ mice during aging. Despite severe thrombocytopenia, TPO^-/-^ and MPL^-/-^ mice did not develop marrow failure during a 2-year follow-up. Marrow and splenic HSCs exhibited different responses to TPO/MPL ablation and exogenous TPO treatment. Splenic niche cells compensated for marrow HSC loss in TPO^-/-^ and MPL^-/-^ mice by upregulating CXCL12 levels. These findings provide new insights into the complex regulation of HSCs by TPO/MPL and reveal a previously unknown link between TPO and CXCL12, two key growth factors for HSC maintenance. Understanding the distinct regulatory mechanisms between marrow and spleen hematopoiesis will help develop novel therapeutic approaches for hematopoietic disorders.

## INTRODUCTION

Thrombopoietin (TPO) and its receptor MPL are key regulators of hematopoietic stem cell (HSC) function(Sitnicka et al. 1996; Qian et al. 2007; Yoshihara et al. 2007) and megakaryocyte/platelet production(Kaushansky et al. 1994; de Sauvage et al. 1994; Kaushansky et al. 1995). Loss-of-function mutations in MPL or TPO cause congenital amegakaryocytic thrombocytopenia in humans, a rare condition characterized by severe thrombocytopenia at birth and often progresses to bone marrow failure within the first decade of life(Geddis 2011; King et al. 2005). Mice lacking TPO or MPL also exhibit marked thrombocytopenia and severely reduced HSC numbers. While previous studies have suggested that TPO/MPL signaling is essential for HSC maintenance(Kimura et al. 1998; Qian et al. 2007; Yoshihara et al. 2007; Sitnicka et al. 1996; Decker et al. 2018), mice lacking TPO/MPL were shown to maintain their blood cell counts (with the exception of thrombocytopenia) and display a normal survival up to one year of age(Alexander et al. 1996; de Sauvage et al. 1996; Qian et al. 2007; Yoshihara et al. 2007). It remains unclear whether these mice will eventually develop marrow failure with longer follow-up.

HSCs reside in a specialized microenvironment known as the hematopoietic niche, which provides signals necessary for their survival, self-renewal, and differentiation. While most hematopoiesis occurs in the bone marrow of adults, HSCs can also be found in extramedullary tissues such as the spleen, lung, and liver, where they contribute to hematopoiesis during times of stress and diseases. In mice and humans, the splenic red pulp is a significant site of extramedullary hematopoiesis. Despite this, the regulation of splenic HSCs is not fully understood, particularly how splenic hematopoiesis responds to TPO/MPL signaling (or its ablation).

In this study, we investigated the effects of TPO/MPL ablation on marrow and splenic hematopoiesis during aging of the TPO knock-out (TPO^-/-^) mice(de Sauvage et al. 1996) and MPL knockout (MPL^-/-^) mice(Alexander et al. 1996). We found that the mice maintained a normal life expectancy compared to wild-type control mice, with no evidence of developing aplastic anemia or marrow failure during 2 years of follow-up. Our study revealed that marrow and splenic HSCs respond differently to both TPO/MPL ablation and exogenous TPO treatment, and the splenic niche cells can compensate for marrow HSC loss in TPO^-/-^ and MPL^-/-^ mice by upregulating their CXCL12 levels. These findings not only provide new insights into the complex regulation of HSCs by TPO/MPL in different hematopoietic niches, but also reveal a previously unknown link between TPO and CXCL12, two critical growth factors essential for HSC maintenance.

## RESULTS

### Long-term maintenance of severe thrombocytopenia without bone marrow failure in TPO^-/-^ and MPL^-/-^ mice

Congenital amegakaryocytic thrombocytopenia is a rare condition characterized by severe thrombocytopenia at birth and patients often progress to aplastic anemia (in which the bone marrow is unable to produce enough blood cells) within the first decade of life(Geddis 2011; King et al. 2005). This condition is caused by missense or non-sense mutations in either the MPL or TPO gene. Previous studies have shown that mice lacking TPO (TPO^-/-^) or MPL (MPL^-/-^) exhibit marked thrombocytopenia and experience a progressive loss of marrow HSCs at 1yr old compared to 3mo old(Qian et al. 2007). Whether TPO^-/-^ and MPL^-/-^ mice will develop multilineage marrow failure during longer follow-up is not known.

To answer this question, we followed the TPO^-/-^ and MPL^-/-^ mice for up to 2 years of age. We found that TPO^-/-^ and MPL^-/-^ mice maintained a severe thrombocytopenia phenotype throughout aging compared to age-matched controls. At 2 years of age, in addition to a severely decreased platelet count (38×10^9^/L in TPO^-/-^ and 107×10^9^/L in MPL^-/-^ vs. 1255×10^9^/L in wild-type controls), TPO^-/-^ and MPL^-/-^ mice developed a moderate decrease in neutrophil count compared to age-matched controls (1.1×10^3^/ul in TPO^-/-^ and 1.7×10^3^/ul in MPL^-/-^ vs. 3.3×10^3^/ul in controls). Hemoglobin and total lymphocyte counts were not significantly different among the groups (Figure 1A). While young (4-6mo old) MPL^-/-^ mice showed a moderate increase in total marrow cell count (1.36-fold) compared to age-matched controls, no significant differences were observed between TPO^-/-^ mice or old MPL^-/-^ mice and age-matched control mice (Figure 1B). Spleen cell count was not significantly different between TPO^-/-^ or MPL^-/-^ mice and age-matched wild-type controls (Figure 1C). There was also no significant difference in overall survival between the TPO^-/-^ or MPL^-/-^ mice and wild-type control mice (Figure 1D). These findings suggest that TPO and its receptor MPL are not essential for HSC maintenance in mice, as the TPO^-/-^ and MPL^-/-^ mice maintained a severe thrombocytopenia phenotype during aging with no evidence of aplastic anemia or marrow failure.

**Figure 1.**
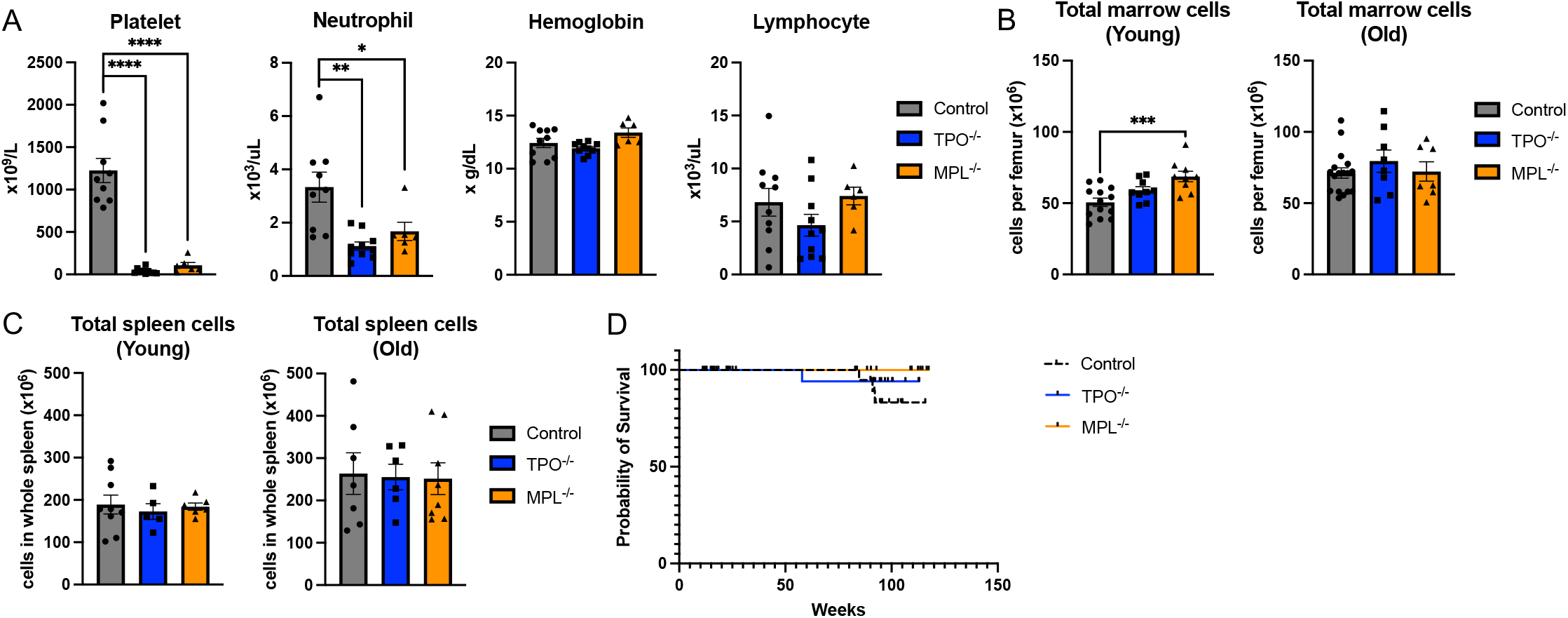
Long-term maintenance of severe thrombocytopenia without bone marrow failure in TPO^-/-^ and MPL^-/-^ mice. (**A**) Peripheral blood cell counts of old (∼24 months old) control *(n=9-10)*, TPO^-/-^ *(n=10)*, and MPL^-/-^ *(n=6)* mice. (**B**) Total marrow cell numbers per femur in young (4-6 months old) (left) and old (right) control *(n=13-17)*, TPO^-/-^ *(n=8-9)*, and MPL^-/-^ *(n=7-9)* mice. (**C**) Total spleen cell numbers in whole spleen in young (left) and old (right) control *(n=7-9)*, TPO^-/-^ *(n=5-6)*, and MPL^-/-^ *(n=6-8)* mice. (**D**) Kaplan-Meier survival curve indicating no significant difference in overall survival between TPO^-/-^ *(n=26)* or MPL^-/-^ *(n=26)* mice and control mice *(n=29). P-values <0*.*05 denoted (*), <0*.*005 denoted (**), <0*.*0005 denoted (***), and <0*.*0001 denoted (****)*.

### Distinct impact of TPO/MPL knockout on marrow and splenic hematopoiesis

The splenic red pulp is known to be a prominent site of extramedullary hematopoiesis during times of stress(Inra et al. 2015). To understand how TPO^-/-^ or MPL^-/-^ mice maintain normal life expectancy and blood cell counts (with the exception of severe thrombocytopenia) despite the absence of TPO and its receptor MPL(Kimura et al. 1998; Qian et al. 2007; Yoshihara et al. 2007; Sitnicka et al. 1996; Decker et al. 2018), we evaluated hematopoiesis in the marrow and spleen of these mice during aging.

We found that marrow Lin^-^cKit^+^Sca1^+^CD150^+^CD48^-^ HSC cell numbers were significantly decreased in young (4-6mo) TPO^-/-^ and MPL^-/-^ mice (4-5-fold) compared to age-matched control mice. This HSC loss was further worsened in old (∼24mo) TPO^-/-^ (25-fold) and MPL^-/-^ (12-fold) mice compared to control mice. However, there was less reduction of Lin^-^cKit^+^Sca1^+^ (LSK) HSPC cells in both young and old TPO^-/-^ and MPL^-/-^ mice (2-3-fold) compared to control mice (Figure 2A-C). These findings indicate a relatively selective role of TPO/MPL ablation in the maintenance of primitive HSCs, with less effect on progenitor cells. In contrast to the bone marrow, there was no significant difference in splenic Lin^-^cKit^+^Sca1^+^CD150^+^CD48^-^ HSC or Lin^-^cKit^+^Sca1^+^ HSPC cell numbers between young TPO^-/-^ or MPL^-/-^ mice and age-matched control mice, with only a moderate decrease of HSCs in old TPO^-/-^ and MPL^-/-^ (3-5-fold) mice compared to control mice (Figure 2D-E).

**Figure 2.**
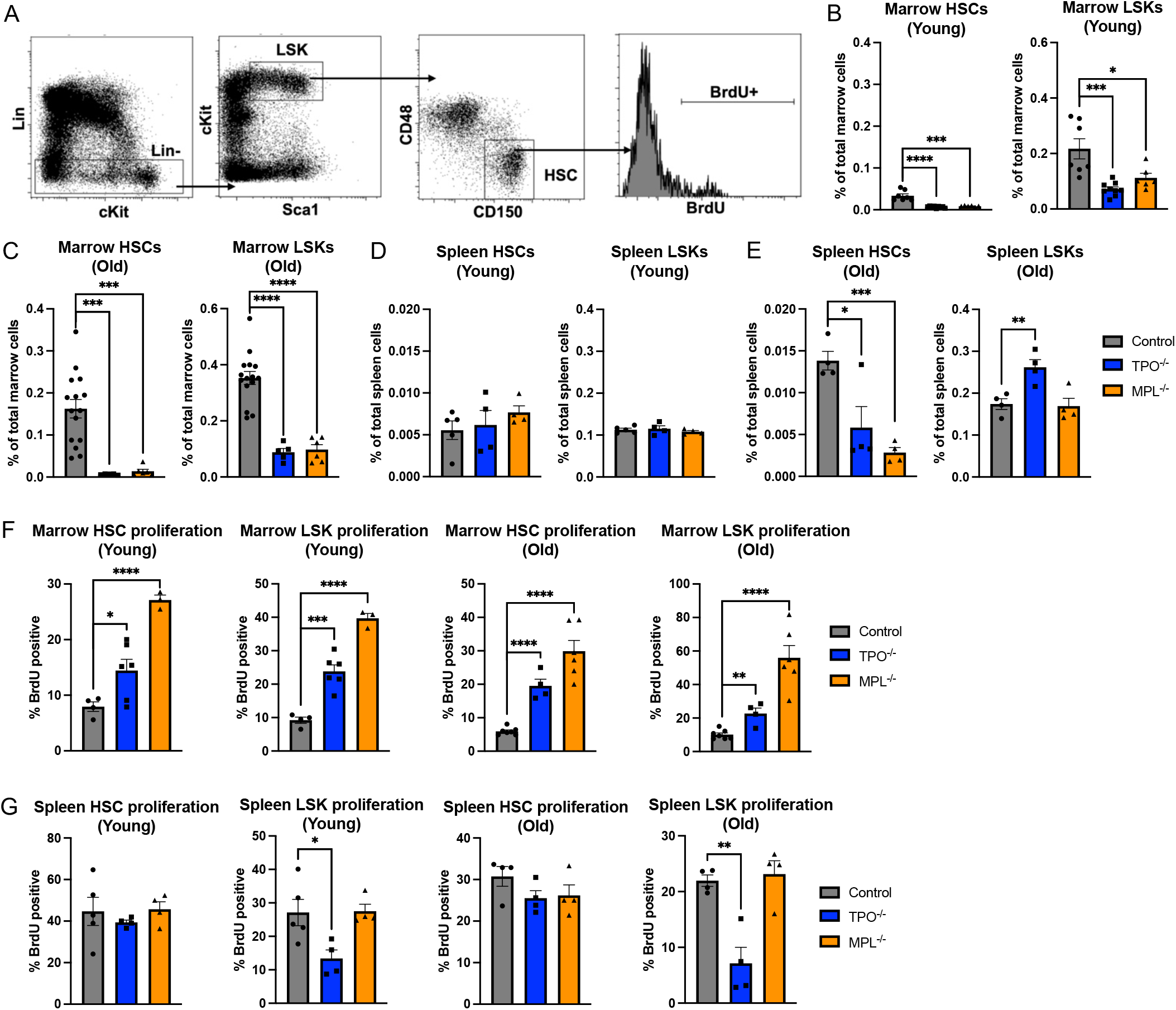
Distinct impact of TPO/MPL knockout on marrow and splenic hematopoiesis. (**A**) Representative flow cytometry plots showing the gating strategy to measure Lin^-^cKit^+^Sca1^+^CD150^+^CD48^-^ HSC and Lin^-^cKit^+^Sca1^+^ HSPC frequency and proliferation. (**B-C**) Marrow HSC frequency and HSPC frequency of young (B) and old (C) control (*n=7-15*), TPO^-/-^ (*n=5-9*), and MPL^-/-^ *(n=6)* mice. (**D-E**) Splenic HSC and HSPC frequency of young (D) and old (E) control (*n=4-5*), TPO^-/-^ (*n=4*), and MPL^-/-^ *(n=4)* mice. (**F-G**) Marrow (F) and spleen (G) HSC and HSPC proliferation of young and old control (*n=4-7*), TPO^-/-^ (*n=4-6*), and MPL^-/-^ *(n=3-6)* mice measured by *in vivo* BrdU labeling.

Previous studies have demonstrated that TPO plays a critical role in maintaining HSC quiescence(Qian et al. 2007; Yoshihara et al. 2007). To investigate the effects of TPO/MPL ablation on HSC proliferation, we measured cell proliferation *in vivo* using BrdU labeling. BrdU is a synthetic nucleoside that is incorporated into the *de novo* synthesized DNA of proliferating cells. Our results showed that, after 2 days of treatment with BrdU, there was an increased proliferation of bone marrow HSCs and LSKs in TPO^-/-^ and MPL^-/-^ mice compared to age-matched control mice, in both young and old mice (Figure 2F). In contrast, there was no significant increase in splenic HSC proliferation in TPO^-/-^ or MPL^-/-^ mice compared to age-matched control mice. We even observed a decreased splenic LSK cell proliferation in TPO^-/-^ mice compared to control mice (Figure 2G).

Taken together, these findings suggest that TPO/MPL signaling has distinct effects on both the number and the function of HSCs in the bone marrow and spleen. Consistent with previous reports(Qian et al. 2007; Yoshihara et al. 2007), TPO/MPL ablation results in increased proliferation and severe age-progressive loss of HSCs in the marrow. In contrast, splenic HSC proliferation is not increased, and the spleen is able to maintain its HSC population in the absence of TPO/MPL signaling.

### Enhanced engraftment capacity of splenic HSCs in comparison to marrow HSCs in TPO/MPL knockout mice

Quiescence is a crucial property of HSCs, as it is tightly linked to their self-renewal and repopulating ability. To further investigate the impact of TPO/MPL ablation on marrow and splenic HSC function, we conducted two types of competitive repopulation assays. In the first assay, CD45.2 marrow cells from 2yr old TPO^-/-^ or MPL^-/-^ mice were injected intravenously together with 2yr old CD45.1 wild-type competitor marrow cells into lethally irradiated CD45.1 recipients. In the second assay, CD45.2 spleen cells from 2yr old TPO^-/-^ or MPL^-/-^ mice were injected together with 2yr old CD45.1 wild-type competitor spleen cells into lethally irradiated CD45.1 recipients (Figure 3A and 3C). We followed the recipients of the marrow donors for 12 weeks and the recipients of the spleen donors for 12 days after transplantation. Our results showed that HSCs from TPO^-/-^ or MPL^-/-^ marrow exhibited severely impaired reconstitution in both the marrow and spleen compared to age-matched wild-type cells. Conversely, HSCs from TPO^-/-^ or MPL^-/-^ spleen demonstrated significantly higher donor engraftment compared to marrow donors from the same mice (Figure 3B and 3D). Moreover, splenic donors from TPO^-/-^ mice showed similar reconstitution in both the marrow and spleen of lethally irradiated mice as the wild-type competitor splenic donor cells (Figure 3B). These results suggest that splenic HSCs have enhanced repopulating capacity compared to marrow HSCs in TPO/MPL ablated mice.

**Figure 3.**
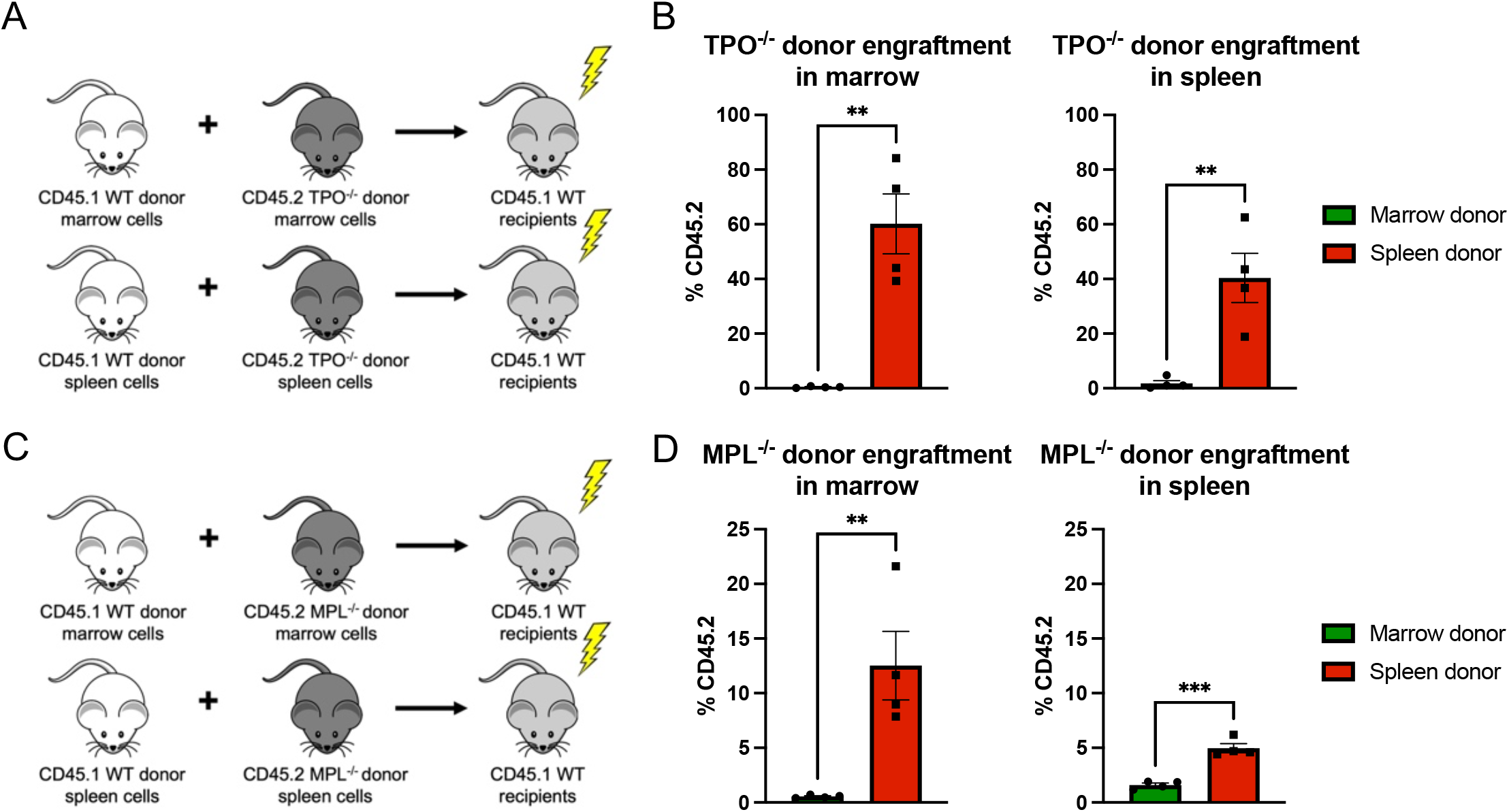
Enhanced engraftment capacity splenic HSCs in comparison to marrow HSCs in TPO^-/-^ and MPL^-/-^ mice. (**A**) Scheme of competitive TPO^-/-^ marrow or splenic donor transplantation experiments. (**B**) Engraftment of CD45.2 TPO^-/-^ marrow and spleen donor cells in the marrow (left) and spleen (right) of wild-type recipient mice (*n=4)*. (**C**) Scheme of competitive MPL^-/-^ marrow or splenic donor transplantation experiments. (**D**) Engraftment of CD45.2 MPL^-/-^ marrow and spleen donor cells in the marrow (left) and spleen (right) of wild-type recipient mice (*n=4)*.

### Distinct gene expression profile of splenic HSCs in comparison to marrow HSCs in TPO/MPL knockout mice

To gain insight into the transcriptomic changes associated with TPO/MPL ablation and enhanced HSC function in the spleen, we performed RNA sequencing on Lineage^-^cKit^+^ HSPCs from the marrow and spleen of wild-type, TPO^-/-^, and MPL^-/-^ mice. We chose to focus on young mice, as both young and old TPO^-/-^ or MPL^-/-^ mice showed similar changes in HSC cell numbers and cell proliferation compared to wild-type control mice (Figure 2). Consistent with the TPO-MPL signaling axis, we found a high degree of similarity between TPO^-/-^ and MPL^-/-^ HSPCs in both marrow and spleen, with Pearson correlation coefficients exceeding 0.90 (Figure 4A). There was a greater degree of gene deregulation between marrow and splenic HSPCs in TPO/MPL knockout mice compared to wild-type mice. Specifically, we identified 2,045 genes that were significantly deregulated between the marrow and splenic HSPCs in wild-type mice, while 3,889 genes were significantly deregulated in TPO/MPL knockout mice. Among these genes, 2,588 were uniquely deregulated in TPO/MPL knockout mice and not in wild-type mice (Figure 4B). These findings indicate that TPO/MPL ablation had a profound impact on splenic hematopoiesis.

**Figure 4.**
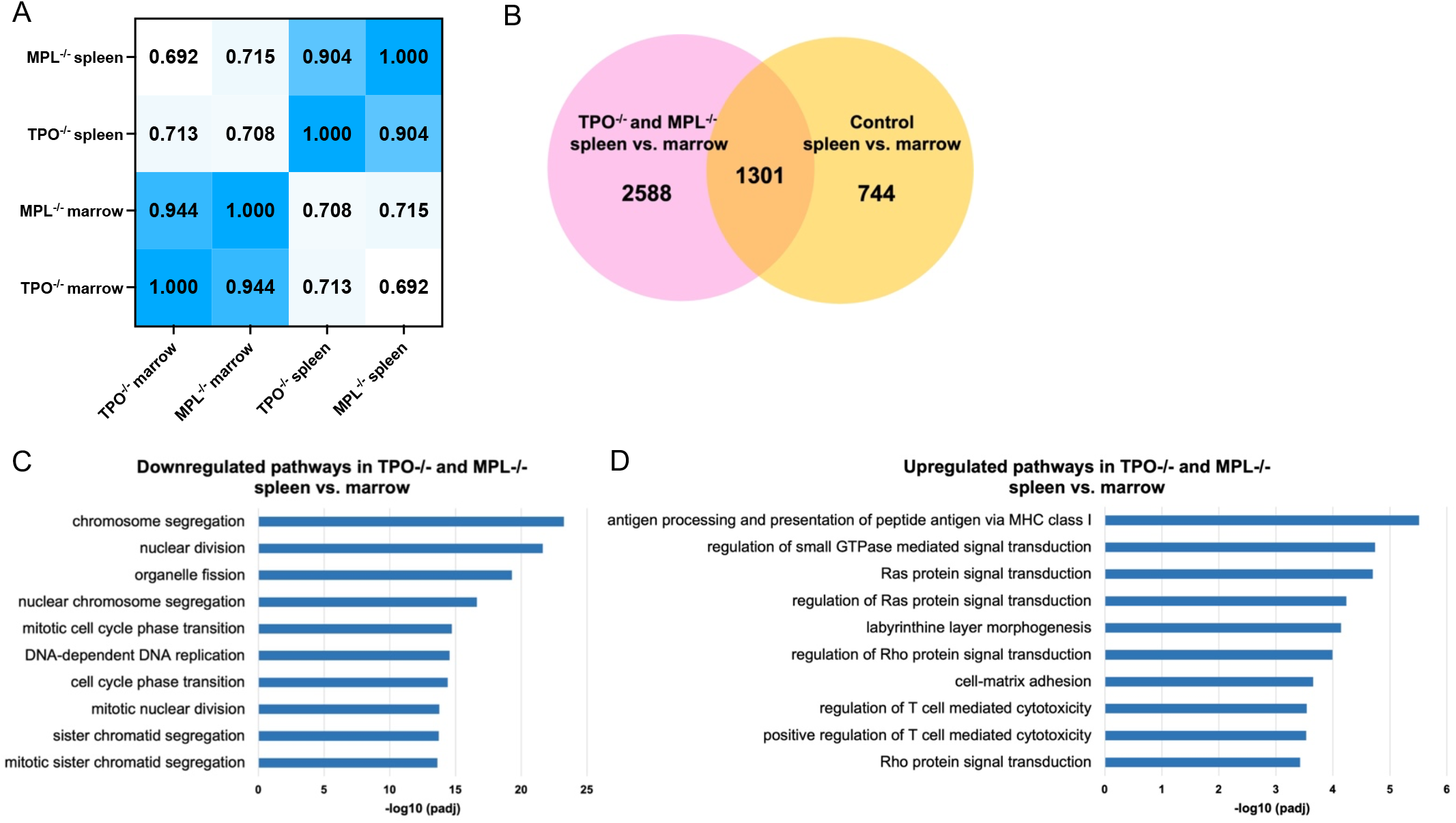
Differential gene expression profile of splenic HSCs in comparison to marrow HSCs in TPO/MPL knockout mice. (**A**) Pearson correlation coefficients for gene expression between TPO^-/-^ and MPL^-/-^ marrow and splenic Lin^-^cKit^+^ HSPCs. TPO^-/-^ and MPL^-/-^ groups consist of a pooled sample of *n=3*. (**B**) Number of deregulated genes in TPO^-/-^ and MPL^-/-^ spleen vs. marrow compared to the number of deregulated genes in control spleen vs. marrow from RNA sequencing analysis. Control groups consists of a pooled sample of *n=2*. (**C-D**) Differentially enriched gene ontology (GO) pathways uniquely downregulated (C) and upregulated (D) in TPO^-/-^ and MPL^-/-^ spleen vs. marrow HSPCs but not in control spleen vs. marrow HSPCs. Adjusted P-values are plotted as the negative of their logarithm.

We examined the gene ontology terms that were exclusively enriched in the comparison between TPO^-/-^ and MPL^-/-^ splenic vs. marrow HSPCs, but not in the comparison between wild-type splenic vs. marrow HSPCs. We found that the most significantly downregulated pathways in TPO^-/-^ and MPL^-/-^ splenic HSPCs were related to cell cycling, DNA replication, and cell division (Figure 4C). This downregulation of cell cycling pathways in TPO^-/-^ and MPL^-/-^ splenic HSPCs supports our observation that TPO/MPL knockout led to increased marrow HSC proliferation, but not splenic HSC proliferation (Figure 2G).

Small GTPase signaling, Ras signaling, Rho signaling, and cell-matrix adhesion pathways were among the most upregulated pathways in TPO^-/-^ and MPL^-/-^ splenic HSPCs compared to marrow HSPCs (Figure 4D). CXCL12 is a crucial niche factor required for maintaining HSC function, including retention, quiescence, and repopulating activity(Tzeng et al. 2011; Ding et al. 2012; Greenbaum et al. 2013; Sugiyama et al. 2006; Peled et al. 1999). CXCL12 binds to its receptor CXCR4, which is a G-protein-coupled-receptor that activates intracellular signaling involving the Ras and Rho small GTPase signaling pathways(Teicher and Fricker 2010). Based on these findings, as well as the maintenance of HSC number, quiescence, and repopulating activity in the spleen but not the marrow of TPO/MPL knockout mice, we hypothesize that the spleen is able to maintain its HSC number and function even in the absence of TPO/MPL signaling by upregulating key hematopoietic niche factors such as CXCL12.

### The impact of TPO/MPL ablation on the bone marrow and splenic hematopoietic niche: a quantitative analysis of niche cells and their expression of niche factors

The adult bone marrow and spleen maintain HSCs through the support of perivascular stromal cells and endothelial cells, which serve as critical sources of factors necessary for HSC maintenance, including CXCL12 and stem cell factor (SCF)(Kiel et al. 2005; Ding et al. 2012; Inra et al. 2015). To investigate how TPO/MPL ablation impacts the hematopoietic niche in both the bone marrow and spleen, we utilized quantitative flow cytometry and/or fluorescence imaging to examine these niche cells and their expression of essential niche factors.

First, we used *in vivo* VE-cadherin labeling and confocal imaging of longitudinally shaved half femurs and thin spleen sections of young wild-type control, TPO^-/-^, and MPL^-/-^ mice to examine marrow and splenic microvasculature. While there was no significant difference in the marrow vascular area (measured by VE-cadherin labeling) between TPO^-/-^ or MPL^-/-^ mice and age-matched control mice, we observed a significant increase in the splenic vascular area in TPO^-/-^ and MPL^-/-^ mice (Figure 5A-B).

**Figure 5.**
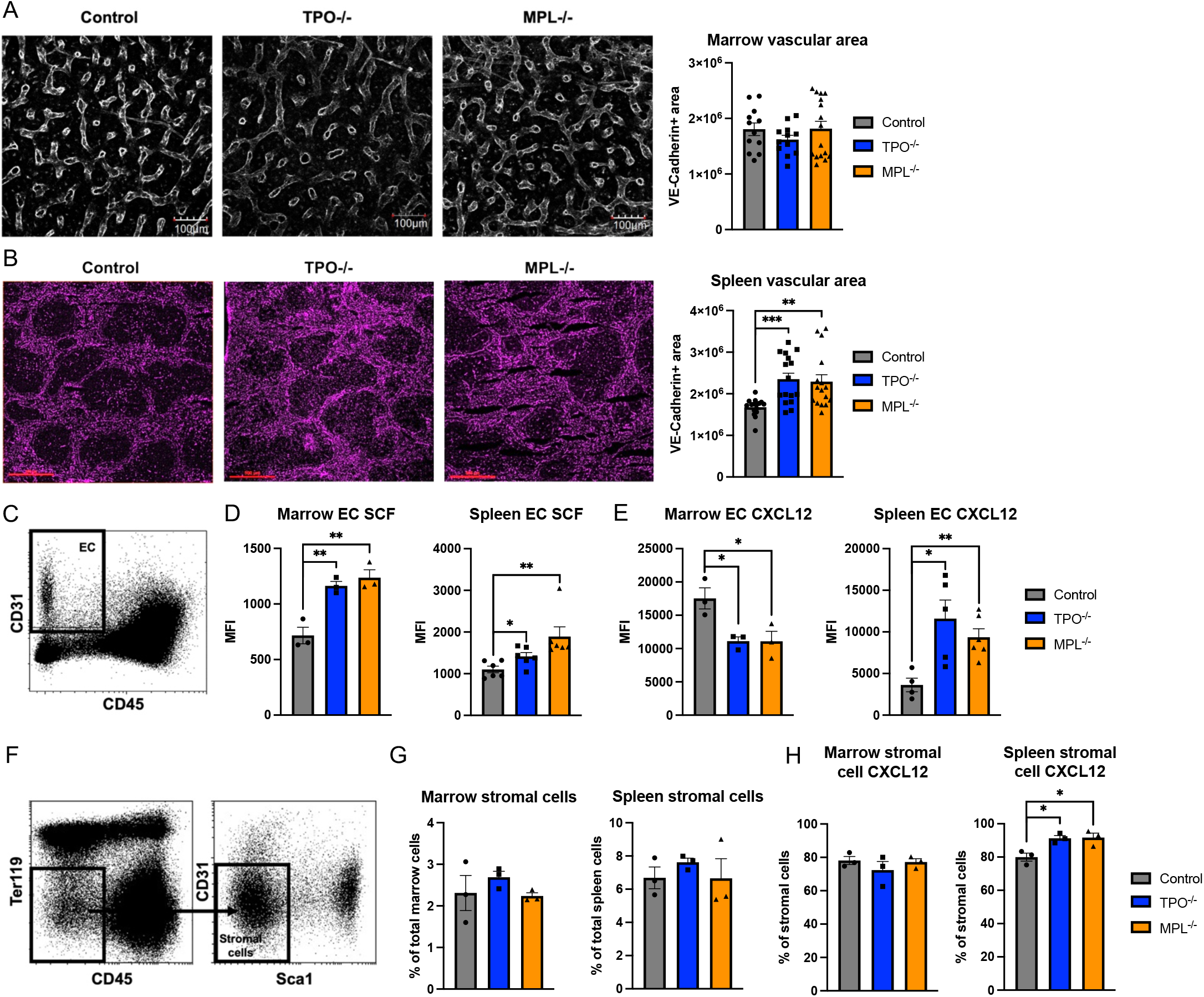
The marrow and splenic hematopoietic niche in TPO^-/-^ and MPL^-/-^ mice. (**A**) Representative whole-mount confocal image of thick femur sections, in which the vasculature was stained intravenously with anti-VE-cadherin antibody (20X magnification, three 5um z-stacks per image, 100um scale bar). Quantification of VE-cadherin^+^ vasculature (white) area in the marrow of young control, TPO^-/-^, and MPL^-/-^ mice (n=2-3 mice in each group). A total of 12-16 non-overlapping 450×350 pixel areas at 20X magnification were analyzed for each group. (**B**) Representative image of thin spleen sections in which the vasculature was stained intravenously with anti-VE-cadherin antibody (4X magnification, 500um scale bar). Quantification of VE-cadherin^+^ vasculature (purple) area in the spleen of young control, TPO^-/-^, and MPL^-/-^ mice (n=2-3 mice in each group). 16 non-overlapping areas (normalized to 450×350 pixels) at 4X were analyzed for each group. (**C-E**) Representative flow cytometry plot of CD45^-^CD31^+^ ECs (C) and flow cytometry quantitative analysis of membranal SCF (D) and intracellular CXCL12 (E) expression in marrow ECs and spleen ECs of young control (n=3-6), TPO^-/-^ (n=3-6), and MPL^-/-^ (n=3-6) mice. MFI: median fluorescence intensity. (**F**) Representative flow cytometry plots showing the gating strategy of CD45^-^CD31^-^Ter119^-^Sca1^-^ perivascular stromal cells. (**G**) Flow cytometry quantitative measurement of perivascular stromal cells in the marrow (left) and spleen (right) of young control (n=3), TPO^-/-^ (n=3), and MPL^-/-^ (n=3) mice. (**H**) CXCL12 expression in marrow (left) and spleen (right) perivascular stromal cells of young control (*n=3*), TPO^-/-^ (*n=3*), and MPL^-/-^ *(n=3)* mice measured using flow cytometry.

Next, we used flow cytometry to analyze the levels of SCF and CXCL12 in bone marrow and splenic CD45^-^ CD31^+^ endothelial cells (Figure 5C). We found a significant increase in SCF levels in both marrow and splenic endothelial cells from TPO^-/-^ and MPL^-/-^ mice compared to control mice (Figure 5D). However, CXCL12 levels were significantly decreased in marrow endothelial cells from TPO^-/-^ and MPL^-/-^ mice compared to age-matched control mice, which is consistent with our previous report(Zhang et al. 2018). Conversely, CXCL12 levels were significantly increased in splenic endothelial cells from TPO^-/-^ and MPL^-/-^ mice (Figure 5E).

We further examined CXCL12 expression in both marrow and splenic perivascular stromal cells (CD45^-^CD31^-^ Ter119^-^Sca1^-^)(Omatsu et al. 2014) using flow cytometry (Figure 5F). The frequencies of marrow or splenic perivascular stromal cells were similar among control, TPO^-/-^ or MPL^-/-^ mice (Figure 5G). CXCL12 expression was moderately increased in splenic perivascular stromal cells from TPO^-/-^ and MPL^-/-^ mice compared to age-matched control mice, while CXCL12 levels remained comparable in marrow perivascular stromal cells between TPO/MPL knockout mice and control mice (Figure 5H). Therefore, TPO/MPL ablation enhanced both the quantity (increased vascular area) and quality (increased CXCL12 levels) of the splenic hematopoietic niche, uniquely contributing to the maintenance of splenic HSC number and function in the absence of TPO/MPL signaling.

### Distinct effects of TPO treatment on marrow and splenic hematopoiesis and niche factors

To further understand the distinct phenotypes observed in the marrow and splenic hematopoiesis from TPO^-/-^ and MPL^-/-^ mice, we administered TPO treatment to wild-type mice (1ug daily intraperitoneal injection for a total of 5-10 days) and evaluated its effects on marrow and splenic hematopoiesis.

After 5 days of TPO treatment, wild-type mice exhibited an increase in spleen weight and spleen cell counts, but a decrease in femur marrow cell counts compared to saline-treated mice (Figure 6A). Flow cytometry analysis revealed that TPO treatment increased bone marrow HSC proliferation but did not increase the number of marrow HSCs (Figure 6B). Conversely, TPO treatment did not alter splenic HSC proliferation but did increase HSC numbers in the spleen (Figure 6C). These results suggest that exogenous TPO administration can mobilize marrow HSCs to the spleen, which is consistent with previous reports(Kuter and Begley 2002). Taken together with the findings from the TPO/MPL knockout mice (Figure 2F-G), our study demonstrates that the TPO/MPL signaling affects splenic and marrow HSCs differently, with the TPO/MPL axis influencing marrow HSC proliferation and quiescence but not splenic HSCs in the same way.

**Figure 6.**
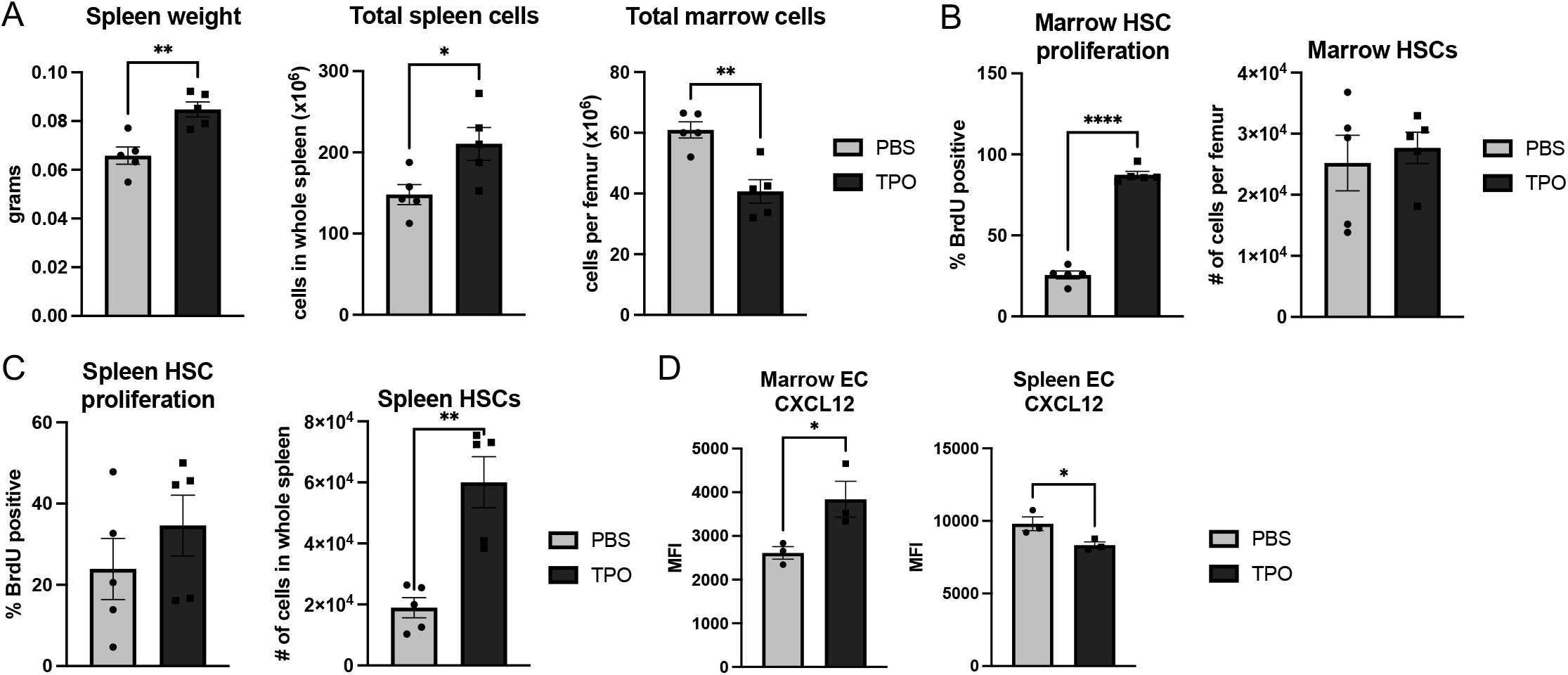
Distinct effects of TPO treatment on marrow and splenic hematopoiesis and niche factors. (**A**) Spleen weight, total spleen cell number, and total marrow cell number per femur of PBS-treated (*n=5*) and TPO-treated (*n=5*) mice. (**B**) Marrow Lin^-^cKit^+^Sca1^+^CD150^+^ CD48^-^ HSC proliferation (left) and frequency (right) of PBS-treated (*n=5*) and TPO-treated (*n=5*) mice. (**C**) Splenic Lin^-^cKit^+^Sca1^+^CD150^+^CD48^-^ HSC frequency (left) and proliferation (right) of PBS-treated (*n=5*) and TPO-treated (*n=5*) mice. (**D**) CXCL12 expression in CD45^-^CD31^+^ ECs from the marrow (left) and spleen (right) of PBS-treated (*n=3*) and TPO-treated (*n=3*) mice.

After administering exogenous TPO for 10 days, we examined CXCL12 expression in the marrow and splenic niche endothelial cells. In contrast to the findings in TPO/MPL knockout mice (Figures 5E), we observed that TPO treatment increased CXCL12 expression in marrow endothelial cells, but decreased CXCL12 levels in splenic endothelial cells (Figure 6D). These observations (Figures 5E and 6D) suggest that TPO can modulate CXCL12 expression, with the opposite effects seen in the marrow and splenic vascular niches.

## DISCUSSIONS

It has long been known that both TPO and MPL are required for HSC maintenance, and loss of TPO/MPL signaling results in the loss of HSC quiescence and the depletion of HSCs(Kimura et al. 1998; Qian et al. 2007; Yoshihara et al. 2007; Sitnicka et al. 1996; Decker et al. 2018). In humans, loss-of-function mutations in MPL or TPO lead to congenital amegakaryocytic thrombocytopenia, which progresses to bone marrow failure within the first decade of life(Geddis 2011; King et al. 2005). However, our findings challenge the notion that TPO/MPL signaling is essential for HSC maintenance in mice, as TPO^-/-^ and MPL^-/-^ mice maintained a severe thrombocytopenia phenotype without developing aplastic anemia or marrow failure during a follow-up period of up to two years. These results highlight the complexity of TPO/MPL signaling in HSC regulation and suggest that the mouse may compensate for TPO/MPL ablation in ways that are not fully understood. A deeper understanding of these compensatory mechanisms could have important implications for the development of novel therapeutic strategies for patients with congenital amegakaryocytic thrombocytopenia.

The spleen is a well-known site of extramedullary hematopoiesis during times of stress in both humans and mice(Inra et al. 2015; Mende et al. 2022). Although the regulation of HSCs likely differs between the marrow and spleen, conclusive evidence is scarce. Our study reveals that TPO/MPL signaling regulates HSC function differently in these two hematopoietic niches by modulating the levels of CXCL12, a crucial niche factor necessary for HSC retention, quiescence, self-renewal, and repopulating activity. Specifically, we found that exogenous TPO administration upregulates CXCL12 expression in marrow niche cells but inhibits CXCL12 expression in splenic niche cells. In contrast, TPO/MPL ablation results in decreased CXCL12 levels, loss of HSC quiescence, and severe age-progressive loss of HSCs in the marrow. However, splenic niche cells can compensate for bone marrow HSC loss in TPO^-/-^ and MPL^-/-^ mice by upregulating their CXCL12 levels, maintaining HSC number and function in the absence of TPO/MPL signaling. Our findings provide mechanistic insights into why TPO/MPL knockout mice maintain a normal life expectancy with no evidence of aplastic anemia or marrow failure. Future research comparing marrow and splenic HSPCs from patients with congenital amegakaryocytic thrombocytopenia will be critical to understand why defective TPO/MPL signaling in these patients leads to marrow failure, in contrast to the findings in TPO^-/-^ and MPL^-/-^ mice.

Our study has also established a novel link between TPO and CXCL12 in the regulation of HSCs. CXCL12 is a crucial niche factor for HSC maintenance in the marrow, and it is primarily produced by endothelial cells and perivascular stromal cells(Ding and Morrison 2013; Greenbaum et al. 2013). However, the mechanisms underlying CXCL12 regulation in the hematopoietic niche remain unclear. Previous studies have reported that CXCL12 can partially restore thrombocytopoiesis in TPO^-/-^ and MPL^-/-^ mice(Avecilla et al. 2004) and TPO administration can promote neovascularization by releasing CXCL12 from platelets(Jin et al. 2006). Our research provides direct evidence that TPO/MPL signaling can modulate HSC function by regulating CXCL12 expression in the vascular niche. Notably, the effects of TPO/MPL on endothelial cell CXCL12 expression are distinct in the bone marrow compared to the spleen. These observations demonstrate that HSC regulation differs between the marrow and spleen, and even the same cytokine (TPO) has unique effects on HSC function in different hematopoietic niches. These findings have significant implications for understanding the regulation of HSCs in different hematopoietic niches.

## EXPERIMENTAL PROCEDURRES

### Experimental mice

TPO knock-out (TPO^-/-^) mice(de Sauvage et al. 1996) were provided by Fred de Sauvage at Genentech (San Francisco, CA) and MPL knockout (MPL^-/-^) mice(Alexander et al. 1996) were provided by Warren Alexander (Melbourne, Australia). All mice used were bred onto a C57BL/6 background and housed and expanded in a pathogen-free mouse facility at Stony Brook University. CD45.1^+^ congenic mice (B6.SJL) were purchased from Taconic Inc (Albany, NY, USA). The experimental groups were not randomized or blinded. All animal experiments were conducted in compliance with the guidelines of the Institutional Animal Care and Use Committee.

### Complete blood counts

Peripheral blood was obtained from the facial vein via submandibular bleeding and collected in EDTA tubes. The samples were then analyzed using a Vetscan Hm5 Hematology Analyzer (Abaxis) to determine the blood count.

### Marrow and spleen cell isolation

Murine femurs and tibias were first harvested and carefully cleaned before marrow cells were flushed out using a 25G needle and syringe into PBS with 2% fetal bovine serum. The remaining bones were then crushed with a mortar and pestle followed by enzymatic digestion with DNase I (25U/ml) and Collagenase D (1mg/ml) at 37 °C for 20 min with gentle rocking. Tissue suspensions were gently mixed using a 10ml pipette to facilitate the dissociation of cellular aggregates. The resulting cell suspensions were then filtered through a 70uM cell strainer to obtain a single-cell suspension.

Murine spleens were collected and placed into a 70uM cell strainer, and then mashed through the strainer into a collecting dish using the plunger end of a 1ml syringe. 10ml PBS with 2% FBS was used to rinse the cell strainer, and the resulting spleen cell suspension was passed through a 10ml syringe with a 23G needle several times to remove any remaining small cell clumps.

### Flow cytometry

All samples were analyzed by flow cytometry using a CytoFLEX flow cytometer (Beckman Coulter, Indianapolis, IN, USA). CD45 (Clone 104) (Biolegend, San Diego, CA, USA), CD45.1 (Clone A20) (BD Biosciences, San Jose, CA), CD45.2 (Clone 104) (Biolegend), Lineage (Lin) cocktail (include CD3, B220, Gr1, CD11b, Ter119; Biolegend), cKit (Clone 2B8, Biolegend), Sca1 (Clone D7, Biolegend), CD150 (Clone mShad150, eBioscience, San Diego, CA, USA), CD48 (Clone HM48-1, Biolegend), CD31 (Clone 390, BD Biosciences), Ter119 (Clone Ter-119, Biolegend) antibodies were used. Intracellular CXCL12 expression and membranal SCF expression were analyzed using an anti-mouse CXCL12 antibody (Clone 79018, R&D Systems, Minneapolis, MN, USA) and an anti-mouse SCF antibody (Clone #40215, R&D Systems) as we previously described(Zhan et al. 2018).

### BrdU incorporation analysis

Mice were injected intraperitoneally with a single dose of 5-bromo-2′-deoxyuridine (BrdU; 100 mg/kg body weight) and maintained on 1 mg BrdU/mL drinking water for 2 days. Mice were then euthanized and marrow and spleen cells isolated as described above. Cells were stained with fluorescent antibodies specific for cell surface markers, followed by fixation and permeabilization using the Cytofix/Cytoperm kit (BD Biosciences), DNase digestion (Sigma, St. Louis, MO), and an anti-BrdU antibody (Clone 3D4, Biolegend) staining to analyze BrdU incorporation as previously described(Castiglione et al. 2021; Lee et al. 2022).

### Competitive transplantation assays

For competitive marrow transplantation, 5 × 10^5^ marrow cells from 2 year old TPO^-/-^ or MPL^-/-^ mice (CD45.2) were injected intravenously together with 5 × 10^5^ marrow cells from age-matched wild-type mice (CD45.1) into lethally irradiated (two 540-rad doses 3 hours apart) 8-week-old wild-type recipient mice (CD45.1).

For competitive spleen transplantation, 1-3 × 10^6^ spleen cells from 2 year old TPO^-/-^ or MPL^-/-^ mice (CD45.2) were injected intravenously together with 1-3 × 10^6^ spleen cells from age-matched wild-type mice (CD45.1) into lethally irradiated (two 540-rad doses 3 hours apart) 8-week-old wild-type recipient mice (CD45.1). Two independent experiments were performed.

### Transcriptome analysis using RNA sequencing

For RNA sequencing experiment, marrow and spleen Lin^-^cKit^+^ hematopoietic stem/progenitor cells (HSPCs) were isolated by magnetic bead isolation (with ∼95% purity) (Miltenyi Biotec) from 3-4mo old wild-type (n=2), TPO^-/-^ (n=3), and MPL^-/-^ (n=3) mice. Total RNA was extracted using the RNeasy mini kit (Qiagen, Hilden, Germany). RNA integrity and quantitation were assessed using the RNA Nano 6000 Assay Kit of the Bioanalyzer 2100 system (Agilent Technologies, CA, USA). For each sample, 400ng of RNA was used to generate sequencing libraries using NEBNextÒ Ultra™ RNA Library Prep Kit for Illumina® (New England BioLabs, MA, USA) following manufacturer’s recommendations. The clustering of the index-coded samples was performed on a cBot Cluster Generation System using PE Cluster Kit cBot-HS (Illumina) according to the manufacturer’s instructions. After cluster generation, the library preparations were sequenced on an Illumina platform. Index of the reference genome was built using hisat2 2.1.0 and paired-end clean reads were aligned to the reference genome using HISAT2. HTSeq v0.6.1 was used to count the reads numbers mapped to each gene. Differential gene expression analysis was performed using the DESeq R package (1.18.0). Genes with an adjusted *p* value < 0.05 found by DESeq were assigned as differentially expressed. Gene Ontology (GO) (http://www.geneontology.org/) and enrichment analysis of differentially expressed genes was implemented by the ClusterProfiler R package. GO terms with corrected p values < 0.05 were considered significantly enriched.

### VE-cadherin in vivo staining and immunofluorescence imaging of marrow and spleen vasculature

Twenty-five micrograms of Alexa Fluor 647-conjugated monoclonal antibody that targets mouse VE-cadherin (clone BV13, Biolegend) was injected retro-orbitally into young and old wild-type, TPO^-/-^, and MPL^-/-^ mice under anesthesia. Ten minutes after antibody injection, the mice were euthanized. Freshly harvested femurs and spleens were fixed in 4% paraformaldehyde in PBS (Affymetrix) for 6 hours at 4 °C while rotating. The tissues were washed in PBS overnight to remove the fixative and cryoprotected in a 20% sucrose PBS solution at 4 °C. The tissues were embedded in OCT (Tissue-Tek) and flash-frozen at -80°C.

To image marrow vasculature, half femur samples were imaged using an Olympus Fluoview FV1000 confocal laser scanning system with a 20X objective magnification, 512 × 512 pixel resolution, and 5 μm *Z*-steps. Half-bone whole-mount tissue sectioning and clearing were performed as described previously(Lee et al. 2022). To image spleen vasculature, 10-μm thick sections were used with a Nikon Eclipse Ts2R Fluorescent Microscope and a 4X objective magnification. Image analysis was performed using the ImageJ software (National Institute of Health), and the VE-cadherin^+^ vascular area was quantified from equal-sized images. The sum of analyzed particles was taken from adjustment of the color threshold, as described previously(Lee et al. 2022).

### Exogenous TPO treatment of wild-type mice

8wk old wild-type (C57B/L6) mice were treated with recombinant human TPO (1ug per day, intraperitoneal injection)(Kaushansky et al. 1994) or PBS for a total of 5 or 10 days.

### Statistical analysis

Statistical analysis was performed using two-tailed Student’s *t* tests with Excel software (Microsoft) and GraphPad Prism software. A *P*-value of <.05 was considered statistically significant. Data are presented as mean ± standard error of the mean (SEM).

### Resource availability

#### Materials and availability

This study did not generate new unique reagents.

#### Data and code availability

The RNA sequencing data generated in this study will be available from the Gene Expression Omnibus.

## ACKNOWLEDGEMENTS

This research was supported by the National Heart, Lung, and Blood Institute grant NIH R01 HL134970 and R01 CA266294 (H.Z.), and VA Merit Award BX003947 and BX005584 (H.Z.). We thank Dr. Kenneth Kaushansky for many fruitful discussions on the TPO/MPL signaling.

## AUTHOR CONTRIBUTION

S.L. performed all the experiments on the project and analyzed the data; H.Z. conceived the projects, designed the experiments, analyzed the data, and interpreted the results; S.L. and H.Z. wrote the manuscript.

